# Genetic factors and age are the strongest predictors of humoral immune responses to common pathogens and vaccines

**DOI:** 10.1101/254706

**Authors:** Petar Scepanovic, Cécile Alanio, Christian Hammer, Flavia Hodel, Jacob Bergstedt, Etienne Patin, Christian W. Thorball, Nimisha Chaturvedi, Bruno Charbit, Laurent Abel, Lluis Quintana-Murci, Darragh Duffy, Matthew L. Albert, Jacques Fellay, The Milieu Intérieur Consortium

**Author notes:** These authors contributed equally to the study (Petar Scepanovic, Cécile Alanio). The *Milieu Intérieur* Consortium is composed of the following team leaders: Laurent Abel (Hôpital Necker), Andres Alcover, Hugues Aschard, Kalla Astrom (Lund University), Philippe Bousso, Pierre Bruhns, Ana Cumano, Caroline Demangel, Ludovic Deriano, James Di Santo, Françoise Dromer, Darragh Duffy, Gérard Eberl, Jost Enninga, Jacques Fellay (EPFL, Lausanne), Magnus Fontes, Antonio Freitas, Odile Gelpi, Ivo Gomperts-Boneca, Serge Hercberg (Université Paris 13), Olivier Lantz (Institut Curie), Claude Leclerc, Hugo Mouquet, Etienne Patin, Sandra Pellegrini, Stanislas Pol (Hôpital Cochin), Antonio Raussel (INSERM UMR 1163 – Institut Imagine), Lars Rogge, Anavaj Sakuntabhai, Olivier Schwartz, Benno Schwikowski, Spencer Shorte, Vassili Soumelis (Institut Curie), Frédéric Tangy, Eric Tartour (Hôpital Européen George Pompidou), Antoine Toubert (Hôpital Saint-Louis), Marie-Noëlle Ungeheuer, Lluis Quintana-Murci^§^, Matthew L. Albert^§^. Additional information can be found at: http://www.pasteur.fr/labex/milieu-interieur. Co-coordinators of the *Milieu Intérieur* Consortium.

## Abstract

**Introduction:** Humoral immune responses to infectious agents or vaccination vary substantially among individuals, and many of the factors responsible for this variability remain to be defined. Current evidence suggests that human genetic variation influences (i) serum immunoglobulin levels, (ii) seroconversion rates, and (iii) intensity of antigen-specific immune responses. Here, we evaluate the impact of intrinsic (age and sex), environmental and genetic factors on the variability of humoral response to common pathogens and vaccines.

**Methods:** We characterized the serological response to 15 antigens from common human pathogens or vaccines, in an age‐ and sex-stratified cohort of 1,000 healthy individuals (*Milieu Intérieur* cohort). Using clinical-grade serological assays, we measured total IgA, IgE, IgG and IgM levels, as well as qualitative (serostatus) and quantitative IgG responses to cytomegalovirus, Epstein-Barr virus, herpes simplex virus 1 & 2, varicella zoster virus, *Helicobacter pylori, Toxoplasma gondii*, influenza A virus, measles, mumps, rubella, and hepatitis B virus. Following genome-wide genotyping of single nucleotide polymorphisms and imputation, we examined associations between ~5 million genetic variants and antibody responses using single marker and gene burden tests.

**Results and discussion:** We identified age and sex as important determinants of humoral response, with older individuals and women having higher rates of seropositivity for most antigens. Genome-wide association studies revealed significant associations between variants in the human leucocyte antigen (HLA) class II region on chromosome 6 and anti-EBV and anti-rubella IgG levels. We used HLA imputation to fine map these associations to amino acid variants in the peptide-binding groove of HLA-DRβ1 and HLA-DPβ1, respectively. We also observed significant associations for total IgA levels with two loci on chromosome 2 and with specific KIR-HLA combinations.

**Conclusions:** Using extensive serological testing and genome-wide association analyses in a well-characterized cohort of healthy individuals, we demonstrate that age, sex and specific human genetic variants contribute to inter-individual variability in humoral response. By highlighting genes and pathways implicated in the normal antibody response to frequently encountered antigens, these findings provide a basis to better understand disease pathogenesis.

## Introduction

Humans are regularly exposed to infectious agents, including common viruses such as cytomegalovirus (CMV), Epstein-Barr virus (EBV) or herpes simplex virus-1 (HSV-1), that have the ability to persist as latent infections throughout life – with possible reactivation events depending on extrinsic and intrinsic factors [1]. Humans also receive multiple vaccinations, which in many cases are expected to achieve lifelong immunity in the form of neutralizing antibodies. In response to each of these stimulations, the immune system mounts a humoral response, triggering the production of specific antibodies that play an essential role in limiting infection and providing long-term protection. Although the intensity of the humoral response to a given stimulation has been shown to be highly variable [2, 3, 4], the genetic and non-genetic determinants of this variability are still largely unknown. The identification of such factors may lead to improved vaccination strategies by optimizing vaccine-induced immunoglobulin G (IgG) protection, or to new understanding of autoimmune diseases, where immunoglobulin levels can correlate with disease severity [5].

Several genetic variants have been identified that account for inter-individual differences in susceptibility to pathogens [6, 7, 8], and in infectious [9] or therapeutic [10] phenotypes. By contrast, relatively few studies have investigated the variability of humoral responses in healthy humans [4, 11, 12]. In particular, Hammer C., *et al.* examined the contribution of genetics to variability in human antibody responses to common viral antigens, and fine-mapped variants at the HLA class II locus that associated with IgG responses. To replicate and extend these findings, we measured IgG responses to 15 antigens from common infectious agents or vaccines as well as total IgG, IgM, IgE and IgA levels in 1,000 well-characterized healthy donors. We used an integrative approach to study the impact of age, sex, non-genetic and genetic factors on humoral responses in healthy humans.

## Methods

### Study participants

The *Milieu Intérieur* cohort consists of 1,000 healthy individuals that were recruited by BioTrial (Rennes, France). The cohort is stratified by sex (500 men, 500 women) and age (200 individuals from each decade of life, between 20 and 70 years of age). Donors were selected based on stringent inclusion and exclusion criteria, previously described [13]. Briefly, recruited individuals had no evidence of any severe/chronic/recurrent medical conditions. The main exclusion criteria were: seropositivity for human immunodeficiency virus (HIV) or hepatitis C virus (HCV); ongoing infection with the hepatitis B virus (HBV) – as evidenced by detectable HBs antigen levels; travel to (sub-)tropical countries within the previous 6 months; recent vaccine administration; and alcohol abuse. To avoid the influence of hormonal fluctuations in women during the peri-menopausal phase, only pre‐ or postmenopausal women were included. To minimize the importance of population substructure on genomic analyses, the study was restricted to self-reported Metropolitan French origin for three generations (*i.e.*, with parents and grandparents born in continental France). Whole blood samples were collected from the 1,000 fasting healthy donors on lithium heparin tubes, from September 2012 to August 2013. The clinical study was approved by the Comité de Protection des Personnes - Ouest 6 on June 13th, 2012, and by the French Agence Nationale de Sécurité du Médicament (ANSM) on June 22nd, 2012. The study is sponsored by Institut Pasteur (Pasteur ID-RCB Number: 2012-A00238-35), and was conducted as a single center study without any investigational product. The protocol is registered under ClinicalTrials.gov (study# NCT01699893).

### Serologies

Total IgG, IgM, IgE, and IgA levels were measured using clinical grade turbidimetric test on AU 400 Olympus at the BioTrial (Rennes, France). Antigen-specific serological tests were performed using clinical-grade assays measuring IgG levels, according to the manufacturer’s instructions. A list and description of the assays is provided in Table S1. Briefly, anti-HBs and anti-HBc IgGs were measured on the Architect automate (CMIA assay, Abbott). Anti-CMV IgGs were measured by CMIA using the CMV IgG kit from Beckman Coulter on the Unicel Dxl 800 Access automate (Beckman Coulter). Anti-Measles, anti-Mumps and anti-Rubella IgGs were measured using the BioPlex 2200 MMRV IgG kit on the BioPlex 2200 analyzer (Bio-Rad). Anti-*Toxoplasma gondi*, and anti-CMV IgGs were measured using the BioPlex 2200 ToRC IgG kit on the BioPlex 2200 analyzer (Bio-Rad). Anti-HSV1 and anti-HSV2 IgGs were measured using the BioPlex 2200 HSV-1 & HSV-2 IgG kit on the BioPlex 2200 analyzer (Bio-Rad). IgGs against *Helicobacter Pylori* were measured by EIA using the PLATELIA *H. Pylori* IgG kit (BioRad) on the VIDAS automate (Biomérieux). Anti-influenza A IgGs were measured by ELISA using the NovaLisa IgG kit from NovaTec (Biomérieux). In all cases, the criteria for serostatus definition (positive, negative or indeterminate) were established by the manufacturer, and are indicated in Table S2. Donors with an unclear result were retested, and assigned a negative result if borderline levels were confirmed with repeat testing.

### Non-genetic variables

A large number of demographical and clinical variables are available in the Milieu Intérieur cohort as a description of the environment of the healthy donors [13]. These include infection and vaccination history, childhood diseases, health-related habits, and sociodemographical variables. Of these, 53 where chosen for subsequent analysis of their impact on serostatus. This selection is based on the one done in [14], with a few variables added, such as measures of lipids and CRP.

### Testing of non-genetic variables

Using serostatus variables as the response, and non-genetic variables as treatment variables, we fitted a logistic regression model for each response and treatment variable pair. A total of 14 * 52 = 742 models where therefore fitted. Age and sex where included as controls for all models, except if that variable was the treatment variable. We tested the impact of the clinical and demographical variables using a likelihood ratio test. All 742 tests where considered a multiple testing family with the false discovery rate (FDR) as error rate.

### Age and sex testing

To examine the impact of age and sex we performed logistic and linear regression analyses for serostatus and IgG levels, respectively. All continuous traits (i.e. quantitative measurements of antibody levels) were log10-transformed in donors assigned as positive using a clinical cutoff. We used false discovery rate (FDR) correction for the number of serologies tested (associations with P < 0.05 were considered significant). To calculate odd ratios in the age analyses, we separated the cohort in equal numbers of young (<45 years old) and old (>=45 years old) individuals, and utilized the epitools R package (v0.5-10).

### DNA genotyping

Blood was collected in 5mL sodium EDTA tubes and was kept at room temperature (18–25°) until processing. DNA was extracted from human whole blood and genotyped at 719,665 single nucleotide polymorphisms (SNPs) using the HumanOmniExpress-24 BeadChip (Illumina). The SNP call rate was higher than 97% in all donors. To increase coverage of rare and potentially functional variation, 966 of the 1,000 donors were also genotyped at 245,766 exonic variants using the HumanExome-12 BeadChip. The HumanExome variant call rate was lower than 97% in 11 donors, which were thus removed from this dataset. We filtered out from both datasets genetic variants that: (i) were unmapped on dbSNP138, (ii) were duplicated, (iii) had a low genotype clustering quality (GenTrain score < 0.35), (iv) had a call rate < 99%, (v) were monomorphic, (vi) were on sex chromosomes, or (vii) diverged significantly from Hardy-Weinberg equilibrium (HWE *P* < 10^−7^). These quality-control filters yielded a total of 661,332 and 87,960 variants for the HumanOmniExpress and HumanExome BeadChips, respectively. Average concordance rate for the 16,753 SNPs shared between the two genotyping platforms was 99.9925%, and individual concordance rates ranged from 99.8% to 100%.

### Genetic relatedness and structure

As detailed elsewhere [14], relatedness was detected using KING [15]. Six pairs of related participants (parent-child, first and second degree siblings) were detected and one individual from each pair, randomly selected, was removed from the genetic analyses. The genetic structure of the study population was estimated using principal component analysis (PCA), implemented in EIGENSTRAT (v6.1.3) [16].

### Genotype imputation

We used Positional Burrows-Wheeler Transform for genotype imputation, starting with the 661,332 quality-controlled SNPs genotyped on the HumanOmniExpress array. Phasing was performed using EAGLE2 (v2.0.5) [17]. As reference panel, we used the haplotypes from the Haplotype Reference Consortium (release 1.1) [18]. After removing SNPs that had an imputation info score < 0.8 we obtained 22,235,661 variants. We then merged the imputed dataset with 87,960 variants directly genotyped on the HumanExome BeadChips array and removed variants that were monomorphic or diverged significantly from Hardy-Weinberg equilibrium (P < 10^−7^). We obtained a total of 12,058,650 genetic variants to be used in association analyses.

We used SNP2HLA (v1.03) [19] to impute 104 4-digit HLA alleles and 738 amino acid residues (at 315 variable amino acid positions of the HLA class I and II proteins) with a minor allele frequency (MAF) of >1%.

We used KIR*IMP [20] to impute KIR alleles, after haplotype inference on chromosome 19 with SHAPEIT2 (v2.r790) [21]. A total of 19 KIR types were imputed: 17 loci plus two extended haplotype classifications (A vs. B and KIR haplotype). A MAF threshold of 1% was applied, leaving 16 KIR alleles for association analysis.

### Genetic association analyses

For single variant association analyses, we only considered SNPs with a MAF of >5% (N=5,699,237). We used PLINK (v1.9) [22] to perform logistic regression for binary phenotypes (serostatus: antibody positive versus negative) and linear regression for continuous traits (log10-transformed quantitative measurements of antibody levels in donors assigned as positive using a clinical cutoff). The first two principal components of a PCA based on genetic data, age and sex were used as covariates in all tests. In order to correct for baseline difference in IgG production in individuals, total IgG levels were included as covariates when examining associations with antigen-specific antibody levels, total IgM, IgE and IgA levels. From a total of 53 additional variables additional co-variates, selected by using elastic net [23] and stability selection [24] as detailed elsewhere [14], were included in some analyses (Table S3). For all antigen-specific genome-wide association studies, we used a genome-wide significant threshold (P_threshold_ < 3.3 x 10^−9^) corrected for the number of antigens tested (N=15). For genome-wide association tests with total Ig levels we set the threshold at P_threshold_ < 1.3 x 10^−8^, correcting for the four immunoglobulin classes tested. For specific HLA analyses, we used PLINK (v1.07) [25] to perform conditional haplotype-based association tests and multivariate omnibus tests at multi-allelic amino acid positions.

### Variant annotation and gene burden testing

We used SnpEff (v4.3g) [26] to annotate all 12,058,650 variants. A total of 84,748 variants were annotated as having (potentially) moderate (e.g. missense variant, inframe deletion, etc.) or high impact (e.g. stop gained, frameshift variant, etc.) and were included in the analysis. We used bedtools v2.26.0 [27] to intersect variant genomic location with gene boundaries, thus obtaining sets of variants per gene. By performing kernel-regression-based association tests with SKAT_CommonRare (testing the combined effect of common and rare variants) and SKATBinary implemented in the SKAT v1.2.1 [28], we tested 16,628 gene sets for association with continuous and binary phenotypes, respectively. By SKAT default parameters, variants with MAF 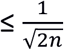 are considered rare, whereas variants with MAF 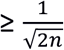 were considered common, where N is the sample size. We used Bonferroni correction for multiple testing, accounting for the number of gene sets and phenotypes tested (P_threshold_ < 2 x 10^−7^ for antigen-specific tests and P_threshold_ < 7.5 x 10^−7^ for tests with total Ig levels).

## Results

### Characterization of humoral immune responses in the 1,000 study participants

To characterize the variability in humoral immune responses between healthy individuals, we measured total IgG, IgM, IgA and IgE levels in the plasma of the 1,000 donors of the *Milieu Interieur* (MI) cohort. After log10 transformation, total IgG, IgM, IgA and IgE levels showed normal distributions, with a median ± sd of 1.02 ±0.08 g/l, 0.01 ±0.2 g/l, 0.31 ±0.18 g/l and 1.51 ±0.62 UI/ml, respectively (Figure S1A).

We then evaluated specific IgG responses to multiple antigens from the following infections and vaccines: (i) 7 common persistent pathogens, including five viruses: CMV, EBV (EA, EBNA, and VCA antigens), herpes simplex virus 1 & 2 (HSV-1 & 2), varicella zoster virus (VZV), one bacterium: *Helicobacter pylori* (H. Pylori), and one parasite: *Toxoplasma gondii* (T. Gondii); (ii) one recurrent virus: influenza A virus (IAV); and (iii) four viruses for which most donors received vaccination: measles, mumps, rubella, and HBV (HBs and HBc antigens) (Figure 1). The distributions of log10-transformed antigen-specific IgG levels in the 1,000 donors for the 15 serologies are shown in Figure S1B. Donors were classified as seropositive or seronegative using the thresholds recommended by the manufacturer (Table S2).

**Fig. 1.**
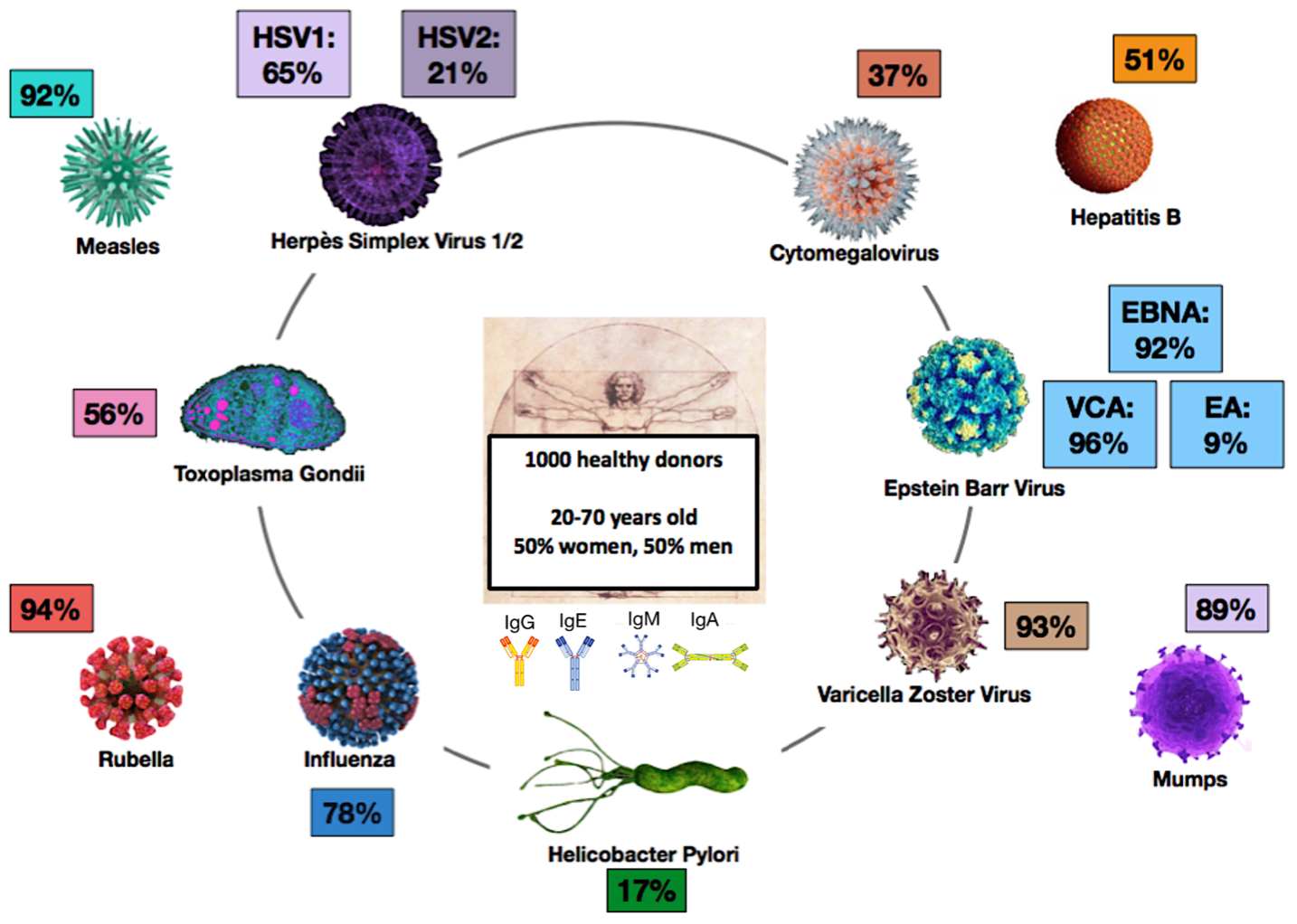
Overview of the study. Serum samples from the 1,000 age‐ and sex-stratified healthy individuals of the *Milieu Intérieur* cohort were used for measuring total antibody levels (IgA, IgM, IgG and IgE), as well as for qualitative (serostatus) and quantitative (IgG levels) assessment of IgG responses against cytomegalovirus, Epstein-Barr virus (anti-EBNA, anti-VCA, anti-EA), herpes simplex virus 1 & 2, varicella zoster virus, *Helicobacter pylori*, *Toxoplasma gondii*, influenza A virus, measles, mumps, rubella, and hepatitis B virus (anti-HBs and anti-HBc), using clinical-grade serological assays.

The vast majority of the 1,000 healthy donors were chronically infected with EBV (seropositivity rates of 96% for EBV VCA, 91% for EBV EBNA and 9% for EBV EA) and VZV (93%). Many also showed high-titer antibodies specific for IAV (77%), HSV-1 (65%), and T. Gondii (56%). By contrast, fewer individuals were seropositive for CMV (35%), HSV-2 (21%), and H. Pylori (18%) (Figure 1, Figure S2A and Table S2). The majority of healthy donors carried antibodies against 5 or more persistent/recurrent infections of the 8 infectious agents tested (Figure S2B). 51% of *MI* donors were positive for anti-HBs IgG - a large majority of them as a result of vaccination, as only 15 study participants (3% of the anti-HBs positive group) were positive for anti-HBc IgG, indicative of previous HBV infection (spontaneously cured, as all donors were negative for HbS antigen, criteria for inclusion in the study). For rubella, measles, and mumps, seropositivity rates were 94%, 91%, and 89% respectively. For the majority of the donors, this likely reflects vaccination with a trivalent vaccine, which was integrated in 1984 as part of national recommendations in France, but for some – in particular the >40 year-old individuals of the cohort, it may reflect acquired immunity due to natural infection.

### Associations of age, sex, and non-genetic variables with serostatus

Subjects included in the *Milieu Interieur* cohort were surveyed for a large number of variables related to infection and vaccination history, childhood diseases, health-related habits, and socio-demographical variables (http://www.milieuinterieur.fr/en/research-activities/cohort/crf-data). Of these, 53 where chosen for subsequent analysis of their impact on serostatus. This selection is based on the one done in [14], with a few variables added, such as measures of lipids and CRP. Applying a mixed model analysis that controls for potential confounders and batch effects, we found expected associations of HBs seropositivity with previous administration of HBV vaccine, as well as of Influenza seropositivity with previous administration of Flu vaccine (Figure S3A and Table S4). We also found associations of HBs seropositivity with previous administration of Typhoid and Hepatitis A vaccines - which likely reflects co-immunization, as well as with Income, Employment, and Owning a house – which likely reflects confounding epidemiological factors.

We observed a significant impact of age on the probability of being seropositive for antigens from persistent or recurrent infectious agents and/or vaccines. For 14 out of the 15 examined serologies, older people (> 45 years old) were more likely to have detectable specific IgG, with an odds ratio (OR; mean ± SD) of 5.4 ± 8.5 (Figure 2A, Figure S3B and Table S5). We identified four different profiles of age-dependent evolution of seropositivity rates (Figure 2B and Figure S4). Profile 1 is typical of childhood-acquired infection, *i.e.* microbes that most donors had encountered by age 20 (EBV, VZV, and influenza). We observed in this case either (i) a limited increase in seropositivity rate after age 20 for EBV; (ii) stability for VZV; or (iii) a small decrease in seropositivity rate with age for IAV (Figure S4A-E). Profile 2 concerns prevalent infectious agents that are acquired throughout life, with steadily increasing prevalence (observed for CMV, HSV-1, and *T. gondii*). We observed in this case either (i) a linear increase in seropositivity rates over the 5 decades of age for CMV (seropositivity rate: 24% in 20-29 years-old; 44% in 60-69 years-old; slope=0.02) and T. Gondii (seropositivity rate: 21% in 20-29 years-old; 88% in 60-69; slope=0.08); or (ii) a nonlinear increase in seropositivity rates for HSV-1, with a steeper slope before age 40 (seropositivity rate: 36% in 20-29 years-old; 85% in 60-69; slope=0.05) (Figure S4F-H). Profile 3 showed microbial agents with limited seroprevalence - in our cohort, HSV-2, HBV (anti-HBS and anti-HBc positive individuals, indicating prior infection rather than vaccination), and *H. Pylori*. We observed a modest increase of seropositivity rates throughout life, likely reflecting continuous low-grade exposure (Figure S4I-K). Profile 4 is negatively correlated with increasing age and is unique to HBV anti-HBs serology (Figure S4L). This reflects the introduction of the HBV vaccine in 1982 and the higher vaccination coverage of younger populations. Profiles for Measles, Mumps and Rubella are provided in Figure S4M-O.

**Fig. 2.**
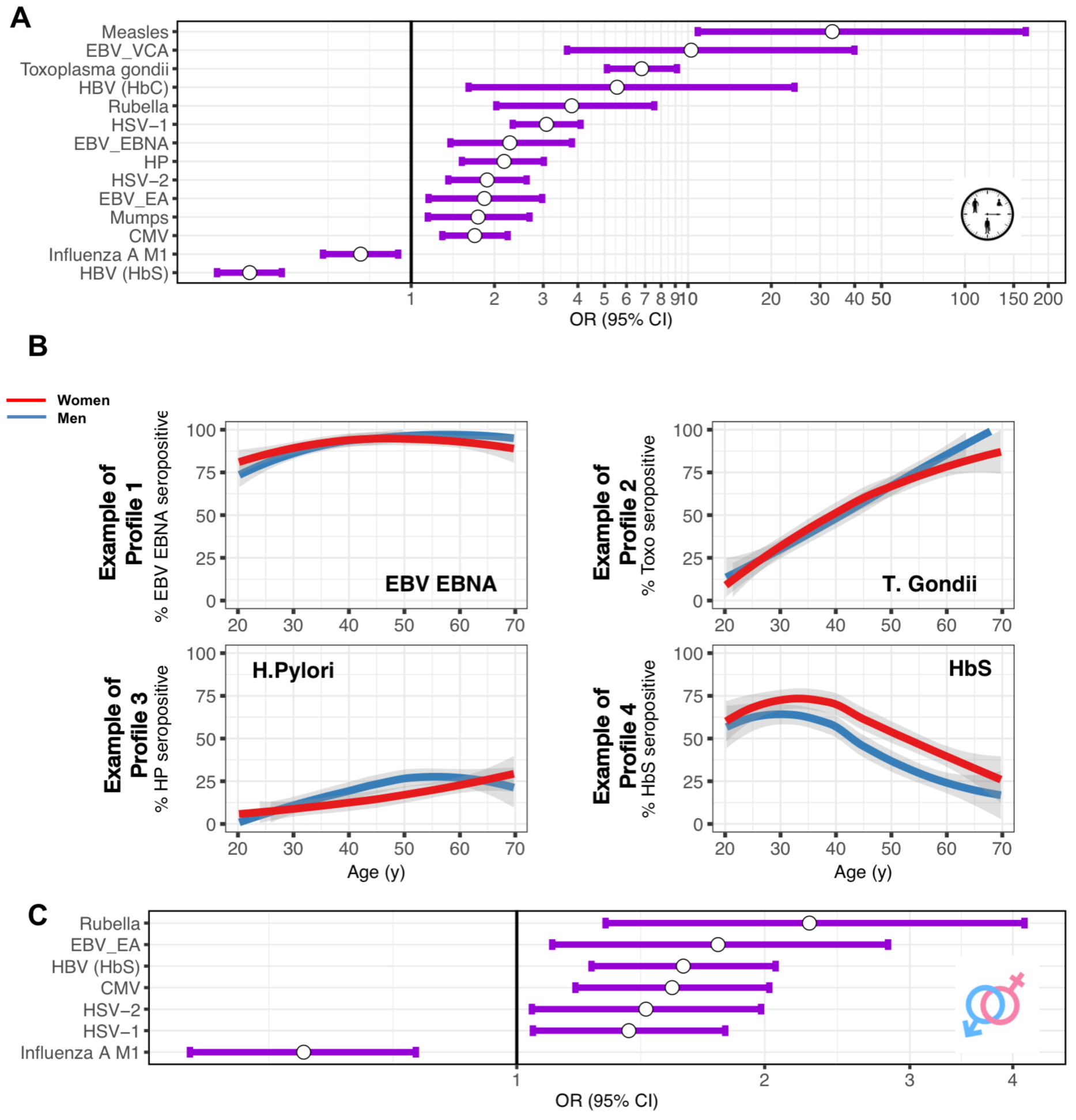
Age and sex impact on serostatus. (**A**) Odd ratios of significant associations (adjusted *P*-values (adj. *P*<0.05) between age (<45 = reference, *vs.* >45 yrs.old) and serostatus as determined based on clinical-grade serologies in the 1,000 healthy individuals from the *Milieu Intérieur* cohort. Odd ratios were estimated in a generalized linear mixed model, with serostatus as response variable, and age and sex as treatment variables. Dots represent the mean of the odd ratios. Lines represent the 95% confidence intervals. (**B**) Odds of being seropositive towards EBV EBNA (Profile 1; upper left), *Toxoplasma gondii* (Profile 2; upper right), *Helicobacter Pylori* (Profile 3; bottom left), and HBs antigen of HBV (Profile 4; bottom right), as a function of age in men (blue) and women (red) in the 1,000 healthy donors. Indicated *P*-values were obtained using a logistic regression with Wald test, with serostatus binary variables (seropositive, versus seronegative) as the response, and age and sex as treatments. (**C**) Odd ratios of significant associations (adjusted *P*-values (adj. *P*<0.05) between sex (Men=reference, *vs.* Women) and serostatus. Odd ratios were estimated in a generalized linear mixed model, with serostatus as response variable, and age and sex as treatment variables. Dots represent the mean of the odd ratios. Lines represent the 95% confidence intervals.

We also observed a significant association between sex and serostatus for 7 of the 15 antigens, with a mean OR of 1.5 ± 0.5 (Figure 2C, Figure S3C and Table S5). For six serological phenotypes, women had a higher rate of positivity, IAV being the notable exception. These associations were confirmed when considering “Sharing house with partner”, and “Sharing house with children” as covariates.

### Impact of age and sex on total and antigen-specific antibody levels

We further examined the impact of age and sex on the levels of total IgG, IgM, IgA and IgE detected in the serum of the patients, as well as on the levels of antigen-specific IgGs in seropositive individuals. We observed a low impact of age and sex with total immunoglobulin levels (Figure 3A and Table S5), and of sex with specific IgG levels (Mumps and VZV; Figure S5A and C). In contrast, age had a strong impact on specific IgG levels in seropositive individuals, affecting 10 out of the 15 examined serologies (Figure 3B, Figure S5B and Table S5). Correlations between age and IgG were mostly positive, *i.e.* older donors had more specific IgG than younger donors, as for example in the case of Rubella (Figure 3C, left panel). The notable exception was *T. gondii*, where we observed lower amounts of specific IgG in older individuals (b=-0.013(−0.019, −0.007), P=3.7x10^−6^, Figure 3C, right panel).

**Fig. 3.**
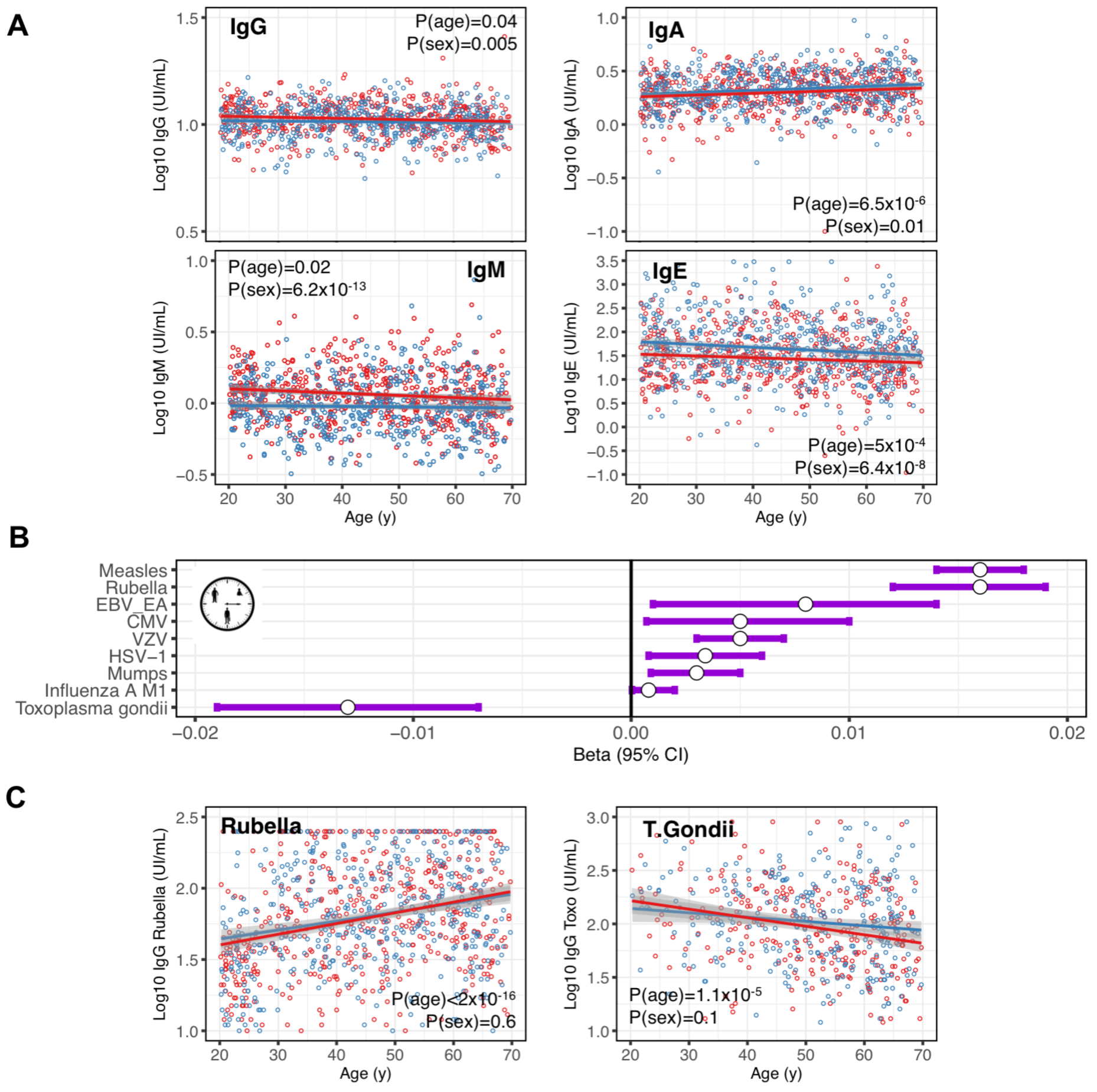
Age and sex impact on total and antigen-specific antibody levels. (**A**) Relationships between Log10-transformed IgG (upper left), IgA (upper right), IgM (bottom left) and IgE (bottom right) levels and age. Regression lines were fitted using linear regression, with Log10-transformed total antibody levels as response variable, and age and sex as treatment variables. Indicated adj. *P* were obtained using the mixed model, and corrected for multiple testing using the FDR method. (**B**) Effect sizes of significant associations (adjusted *P*-values (adj. *P*<0.05) between age and Log10-transformed antigen-specific IgG levels in the 1,000 healthy individuals from the *Milieu Intérieur* cohort. Effect sizes were estimated in a linear mixed model, with Log10-transformed antigen-specific IgG levels as response variables, and age and sex as treatment variables. Dots represent the mean of the beta. Lines represent the 95% confidence intervals. (**C**) Relationships between Log10-transformed anti-rubella IgGs (left), and Log10-transformed anti-toxoplasma gondii IgGs (right) and age. Regression lines were fitted using linear regression described in (**B**). Indicated adj. *P* were obtained using the mixed model, and corrected for multiple testing using the FDR method.

### Genome-wide association study of serostatus

To test if human genetic factors influence the rate of seroconversion upon exposure, we performed genome-wide association studies. Specifically, we searched for associations between 5.7 million common polymorphisms (MAF > 5%) and the 15 serostatus in the 1,000 healthy donors. Based on our results regarding age and sex, we included both as covariates in all models. After correcting for the number of antigens tested, the threshold for genome-wide significance was P_threshold_ = 3.3 x 10^−9^, for which we did not observe any significant association. In particular, we did not replicate the previously reported associations with *H. Pylori* serostatus on chromosome 1 (rs368433, P = 0.67, OR = 0.93) and 4 (rs10004195, P = 0.83, OD = 0.97) [29].

We then focused on the HLA region and confirmed the previously published association of influenza A serostatus with specific amino acid variants of HLA class II molecules [11]. The strongest association in the *MI* cohort was found with residues at position 31 of the HLADRβ1 subunit (omnibus P = 0.009, Table S6). Residues found at that position, isoleucine (P = 0.2, OD (95% CI) = 0.8 (0.56, 1.13)) and phenylalanine (P = 0.2, OR (95% CI) = 0.81 (0.56, 1.13)), are consistent in direction and in almost perfect linkage disequilibrium (LD) with the glutamic acid residue at position 96 in HLA-DRβ1 that was identified in the previous study (Table S7). As such, our result independently validates the previous observation.

### Genome-wide association study of total and antigen-specific antibody levels

To test whether human genetic factors also influence the intensity of antigen-specific immune response, we performed genome-wide association studies of total IgG, IgM, IgA and IgE levels, as well as antigen-specific IgG levels.

Using a significance threshold of P_threshold_ < 1.3 x 10^−8^, we found no SNPs associated with total IgG, IgM, IgE and IgA levels. However, we observed nominal significance and the same direction of the effect for 3 out of 11 loci previously published for total IgA [12, 30, 31, 32, 33], 1 out of 6 loci for total IgG [12, 30, 34] and 4 out of 11 loci for total IgM [12, 35] (Table S8). Finally, we also report a suggestive association (genome-wide significant, P < 5.0 x 10^−8^, but not significant when correcting for the number of immunoglobulin classes tested in the study) of a SNP rs11186609 on chromosome 10 with total IgA levels (P = 2.0 x 10^−8^, beta = −0.07 for the C allele). The closest gene for this signal is *SH2D4B*.

We next explored associations between human genetic variants and antigen-specific IgG levels in seropositive donors (P_threshold_ < 3.3 x 10^−9^). We detected significant associations for anti-EBV (EBNA antigen) and anti-rubella IgGs. Associated variants were in both cases located in the HLA region on chromosome 6. For EBV, the top SNP was rs74951723 (P = 3 x 10^−14^, beta = 0.29 for the A allele) (Figure 4A). For rubella, the top SNP was rs115118356 (P = 7.7 x 10^−10^, beta = −0.11 for the G allele) (Figure 4B). rs115118356 is in LD with rs2064479, which has been previously reported as associated with titers of anti-rubella IgGs (r^2^ = 0.53 and D’ = 0.76) [36].

**Fig. 4.**
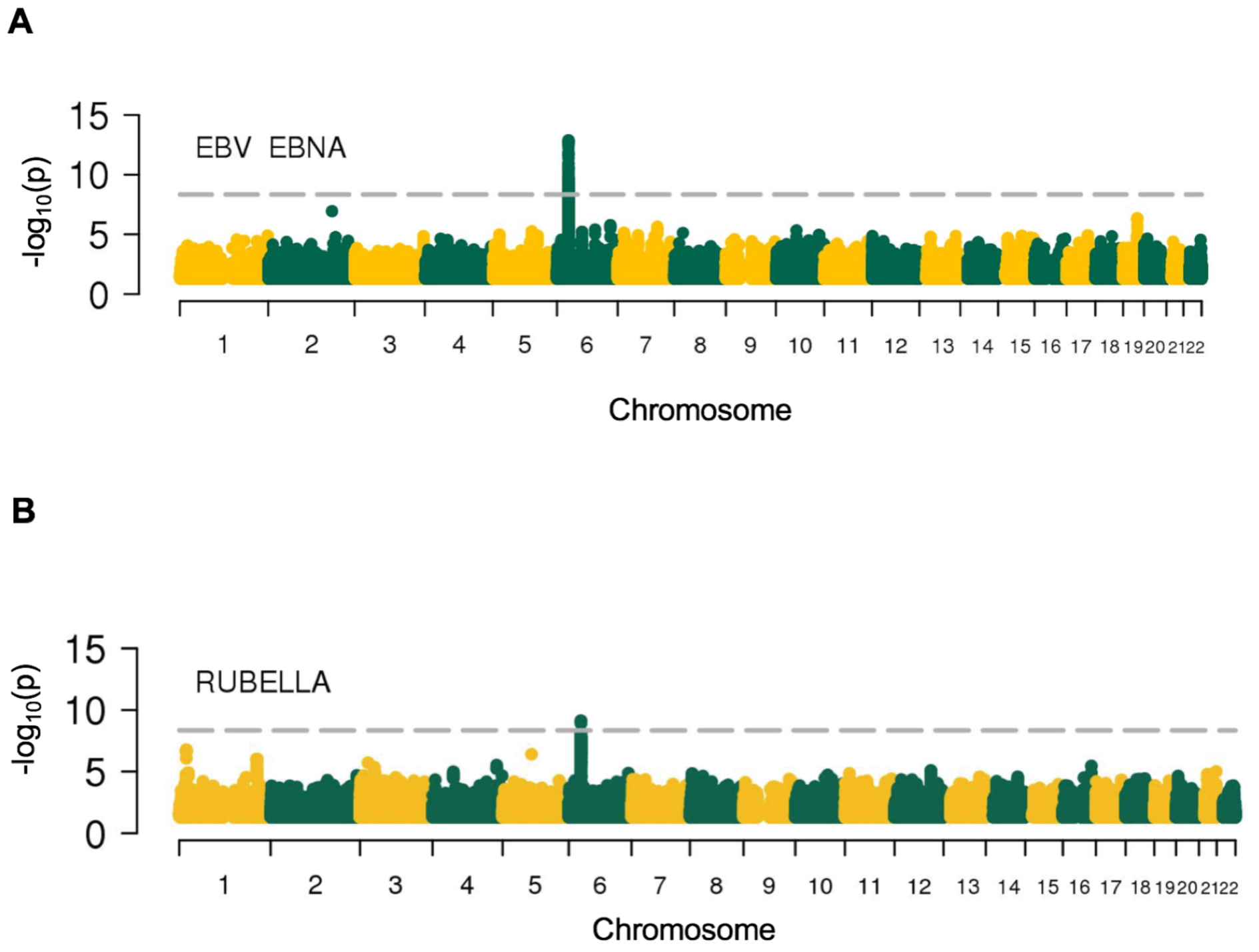
Association between host genetic variants and serological phenotypes. Manhattan plots of association results for (A) EBV anti-EBNA IgG, (B) Rubella IgG levels. The dashed horizontal line denotes genome-wide significance (P = 3.3 x 10^−9^).

To fine map the associations observed in the HLA region, we tested 4-digit HLA alleles and variable amino positions in HLA proteins. At the level of HLA alleles, *HLA-DQB1*03:01* showed the lowest P-value for association with EBV EBNA (P = 1.3 x 10^−7^), and *HLADPB1*03:01* was the top signal for rubella (P = 3.8 x 10^−6^). At the level of amino acid positions, position 58 of the HLA-DR 1 protein associated with anti-EBV (EBNA antigen) IgG levels (P = 2.5 x 10^−11^). This is consistent with results of previous studies linking genetic variations in HLA-DRβ1 with levels of anti-EBV EBNA-specific IgGs [4, 11, 37] (Table S9). In addition, position 8 of the HLA-DP 1 protein associated with anti-rubella IgG levels (P = 1.1 x 10^−9^, Table 1). Conditional analyses on these amino-acid positions did not reveal any additional independent signals.

**Table 1.** Significant associations with EBV EBNA and Rubella antigens at the level of SNP, HLA allele and protein amino acid position

|  |  | Phenotype |  |
| --- | --- | --- | --- |
|  |  | EBV EBNA IgG levels | Rubella IgG levels |
| SNP | ID (Allele) | rs74951723 (A) | rs115118356 (G) |
|  | P-value | 3 x 10<sup>-14</sup> | 7.68 x 10<sup>-10</sup> |
|  | Beta (95% CI) | 0.29 (0.21, 0.36) | -0.11 (-0.15, -0.08) |
| Classical HLA allele | Allele | HLA-DQB1*03:01 | HLA-DPB1*03:01 |
|  | P-value | 1.26 x 10<sup>-7</sup> | 3.8 x 10<sup>-6</sup> |
|  | Beta (95% CI) | 0.17 (0.11, 0.23) | -0.12 (-0.18, -0.07) |
| Amino acid | Protein (position) | HLA-DRβ1 (56) | HLA-DPβ1 (8) |
|  | Omnibus P-value | 2.53 x 10<sup>-11</sup> | 1.12 x 10<sup>-9</sup> |

### KIR associations

To test whether specific KIR genotypes, and their interaction with HLA molecules, are associated with humoral immune responses, we imputed KIR alleles from SNP genotypes using KIR*IMP. First, we searched for potential associations with serostatus or IgG levels for 16 KIR alleles that had a MAF > 1%. After correction for multiple testing, we did not find any significant association (P_threshold_ < 2.6 x 10^−4^). Second, we tested specific KIR-HLA combinations. We filtered out rare combinations by removing pairs that were observed less then 4 times in the cohort. After correction for multiple testing (P_threshold_ < 5.4 × 10^−7^), we observed significant associations between total IgA levels and the two following HLA-KIR combinations: HLA-B*14:02/ KIR3DL1 and HLA-C*08:02/ KIR2DS4 (P = 3.9 x 10^−9^ and P = 4.9 x 10^−9^ respectively, Table 2).

**Table 2.** Association testing between KIR-HLA interactions and serology phenotypes

| Phenotype | KIR | HLA | Estimate | Std. Error | P-value |
| --- | --- | --- | --- | --- | --- |
| IgA levels | KIR3DL1 | HLA-B*14:02 | 0.456 | 0.077 | 3.9x10<sup>-09</sup> |
| IgA levels | KIR2DS4 | HLA-B*14:02 | 0.454 | 0.077 | 4.5x10<sup>-09</sup> |
| IgA levels | KIR3DL1 | HLA-C*08:02 | 0.449 | 0.076 | 4.9x10<sup>-09</sup> |
| IgA levels | KIR2DS4 | HLA-C*08:02 | 0.448 | 0.076 | 5.7x10<sup>-09</sup> |

### Burden testing for rare variants

Finally, to search for potential associations between the burden of low frequency variants and the serological phenotypes, we conducted a rare variant association study. This analysis only included variants annotated as missense or putative loss-of-function (nonsense, essential splice-site and frame-shift, N=84,748), which we collapsed by gene and tested together using the kernel-regression-based association test SKAT [28]. We restricted our analysis to genes that contained at least 5 variants. Two genes were identified as significantly associated with total IgA levels using this approach: *ACADL* (P = 3.4 x 10^−11^) and *TMEM131* (P=7.8 x 10^−11^) (Table 3). By contrast, we did not observe any significant associations between rare variant burden and antigen-specific IgG levels or serostatus.

**Table 3.** Significant associations of rare variants collapsed per gene set with IgA levels.

| Phenotype | Chromosome | Gene | P-value | Q | N° of Rare Markers | N° of Common Markers |
| --- | --- | --- | --- | --- | --- | --- |
| IgA levels | 2 | ACADL | $3.42 \times 10^{-11}$ | 18.09 | 5 | 2 |
| | 2 | TMEM131 | $7.83 \times 10^{-11}$ | 17.89 | 13 | 2 |

## Discussion

We performed genome-wide association studies for a number of serological phenotypes in a well-characterized age‐ and sex-stratified cohort, and included a unique examination of genetic variation at HLA and KIR loci, as well as KIR-HLA associations. As such, our study provides a broad resource for exploring the variability in humoral immune responses across different isotypes and different antigens in humans.

Using a fine-mapping approach, we replicated the previously reported associations of variation in the HLA-DR 1 protein with influenza A serostatus and anti-EBV IgG titers [4, 11], implicating amino acid residues in strong LD with the ones previously reported (Hammer et al.). We also replicated an association between HLA class II variation and anti-Rubella IgG titers [36], and further fine-mapped it to position 8 of the HLA-DP 1 protein. Interestingly, position 8 of HLA-DP 1, as well as positions 58 and 31 of HLA-DR 1, are all part of the extracellular domain of the respective proteins. Our findings confirm these proteins as critical elements for the presentation of processed peptide to CD4^+^ T cells, and as such may reveal important clues in the fine regulation of class II antigen presentation. We also identified specific HLA/KIR combinations, namely HLA-B*14:02/KIR3DL1 and HLAC*08:02/KIR2DS4, which associate with higher levels of circulating IgA. Given the novelty of KIR imputation method and the lack of possibility of benchmarking its reliability in the *MI* cohort further replication of these results will be needed. Yet these findings support the concept that variations in the sequence of HLA Class II molecules, or specific KIRs/HLA class I interactions play a critical role in shaping humoral immune responses in humans. In particular, our findings confirm that small differences in the capacity of HLA class II molecules to bind specific viral peptides can have a measurable impact on downstream antibody production. As such, our study emphasizes the importance of considering HLA diversity in disease association studies where associations between IgG levels and autoimmune diseases are being explored.

We identified nominal significance for some but not all of the previously reported associations with levels of total IgG, IgM and IgA, as well as a suggestive association of total IgA levels with an intergenic region on chromosome 10 - closest gene being *SH2D4B.* By collapsing the rare variants present in our dataset into gene sets and testing them for association with the immunoglobulin phenotypes, we identified two additional loci that participate to natural variation in IgA levels. These associations mapped to the genes *ACADL* and *TMEM131*. *ACADL* encodes an enzyme with long-chain acyl-CoA dehydrogenase activity, and polymorphisms have been associated with pulmonary surfactant dysfunction [38]. As the same gene is associated with levels of circulating IgA in our cohort, we speculate that *ACADL* could play a role in regulating the balance between mucosal and circulating IgA. Further studies will be needed to test this hypothesis, as well as the potential impact of our findings in other IgA-related diseases.

We were not able to replicate previous associations of *TLR1* and *FCGR2A* locus with serostatus for *H. Pylori* [29]. We believe this may be a result of notable differences in previous exposure among the different cohorts as illustrated by the different levels of seropositivity; 17% in the *Milieu Interieur* cohort, versus 56% in the previous ones, reducing the likelihood of replication due to decreased statistical power.

In addition to genetics findings, our study re-examined the impact of age and sex, as well as non-genetic variables, on humoral immune responses. Although this question has been previously addressed, our well-stratified cohort brings interesting additional insights. One interesting finding is the high rate of seroconversion for CMV, HSV-1, and T. Gondii during adulthood. In our cohort, the likelihood of being seropositive for one of these infections is comparable at age 20 and 40. Given the high prevalence of these microbes in the environment, this raises questions about the factors that prevent some individuals from becoming seropositive upon late life exposure.Second, both age and sex have a strong correlation with serostatus, *i.e.* older and female donors were more likely to be seropositive. Although increased seropositivity with age probably reflects continuous exposure, the sex effect is intriguing. Indeed, our study considered humoral responses to microbial agents that differ significantly in terms of physiopathology and that do not necessarily have a childhood reservoir. Also, our analysis show that associations persist after removal of potential confounding factors such as marital status, and/or number of kids. As such, we believe that our results may highlight a general impact of sex on humoral immune response variability, *i.e.* a tendency for women to be more likely to seroconvert after exposure, as compared to men of same age. This result is in line with observations from vaccination studies, where women responded to lower vaccine doses [39]. Finally, we observed an age-related increase in antigen-specific IgG levels in seropositive individuals for most serologies, with the notable exception of toxoplasmosis. This may indicate that aging plays a general role in IgG production. An alternative explanation that requires further study is that this could be the consequence of reactivation or recurrent exposure.

In sum, our study provides evidence that age, sex and host genetics contribute to natural variation in humoral responses in humans. The identified associations have the potential to help improve vaccination strategies, and/or dissect pathogenic mechanisms implicated in human diseases related to immunoglobulin production such as autoimmunity.

## Acknowledgements

This work benefited from support of the French government’s Invest in the Future Program, managed by the Agence Nationale de la Recherche (ANR, reference 10-LABX-69-01). It was also supported by a grant from the Swiss National Science Foundation (31003A_175603, to JF). C.A. received a PostDoctoral Fellowship from Institut National de la Recherche Médicale.

---

**Fig. S1.**
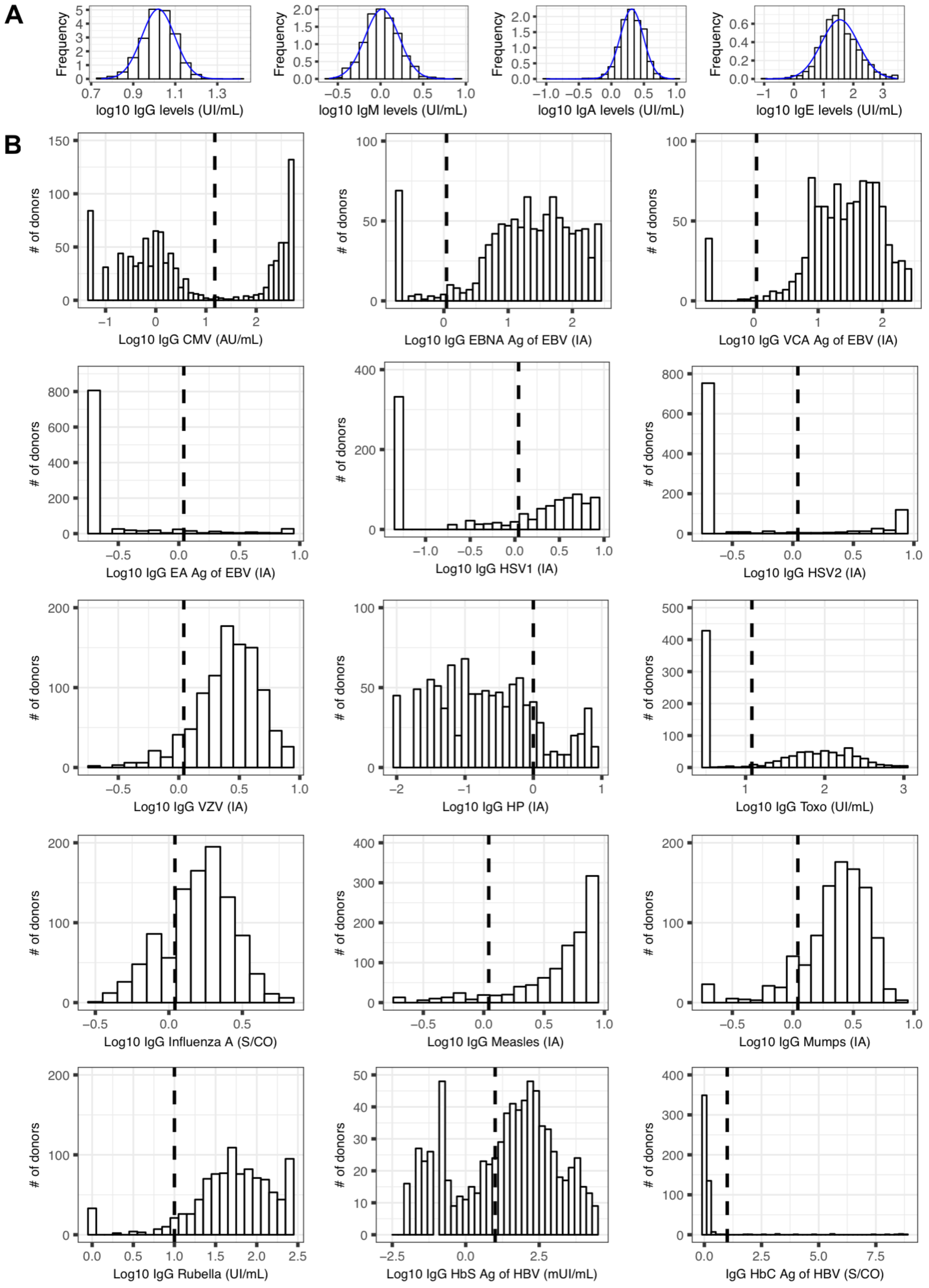
Distribution of serological variables, and clinical threshold used for determination of serostatus. (**A**) Distribution and probability density curve of Log10-transformed IgG, IgM, IgA, IgE levels in the 1,000 study participants. (**B**) Distribution of Log10-transformed antigen-specific IgG levels. The vertical lines indicate the clinical threshold determined by manufacturer, and used for determining the serostatus of the donors for each serology.

**Fig. S2.**
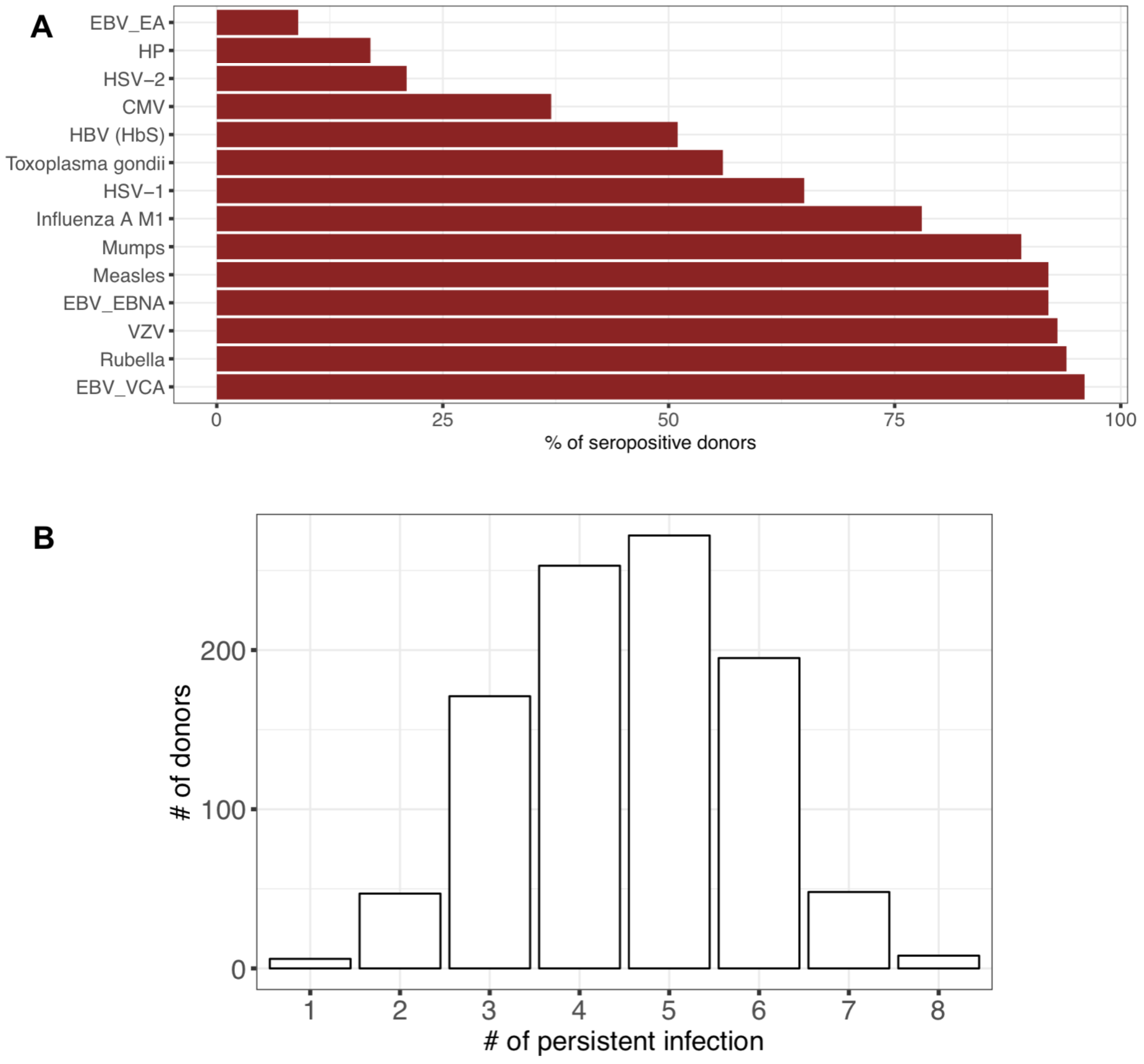
Seroprevalence data in the 1,000 healthy donors. (**A**) Percentage of seropositive donors for each indicated serology in the *MI* study (for HBV serology, percentages of anti-HBs IgGs are indicated). (**B**) Distribution of the number of positive serologies in the 1,000 healthy donors regarding the 8 persistent or recurrent infections tested in our study (*i.e.* CMV, Influenza, HSV1, HSV2, TP, EBV_EBNA, VZV, HP).

**Fig. S3.**
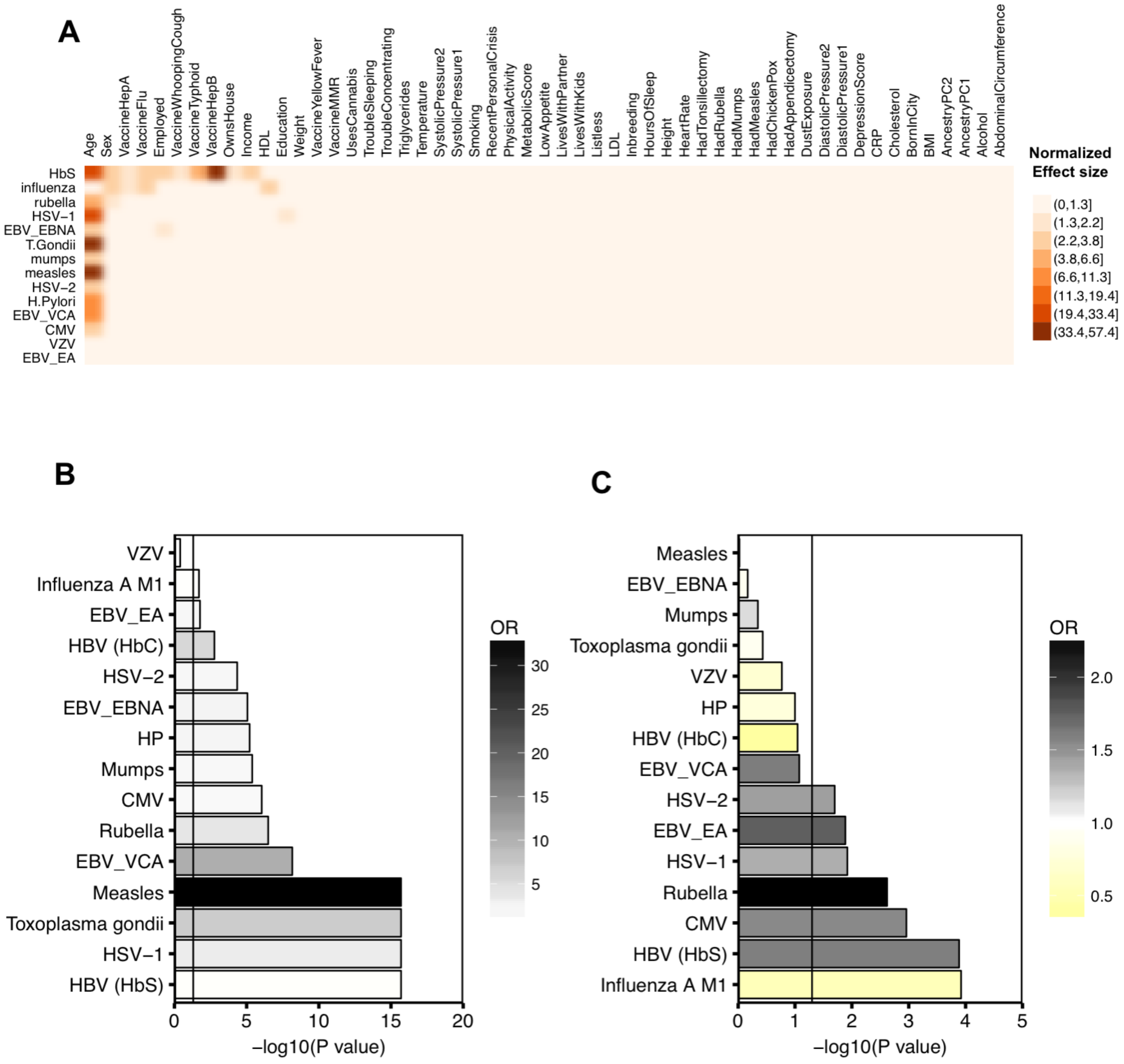
Impact of non-genetic factors, age, and sex on serostatus. (**A**) Adjusted *P*-values (FDR) of the large-sample chi-square likelihood ratio tests of effect of non-genetic variables on serostatus, obtained from mixed models. (**B-C**) Adjusted *P*-values (adj. *P*) of the tests of effect of age (<45 = reference, *vs.* >45 years old) (**B**) and sex (Men = reference, *vs.* Women) (**C**) on serostatus, obtained using a generalized linear mixed model, with serostatus as response variables, and age and sex as treatment variables. Odd ratios were color-coded. Vertical black line indicates the ‐log10 of the chosen threshold for statistical significance (-log10(0.05) = 1.30103).

**Fig. S4.**
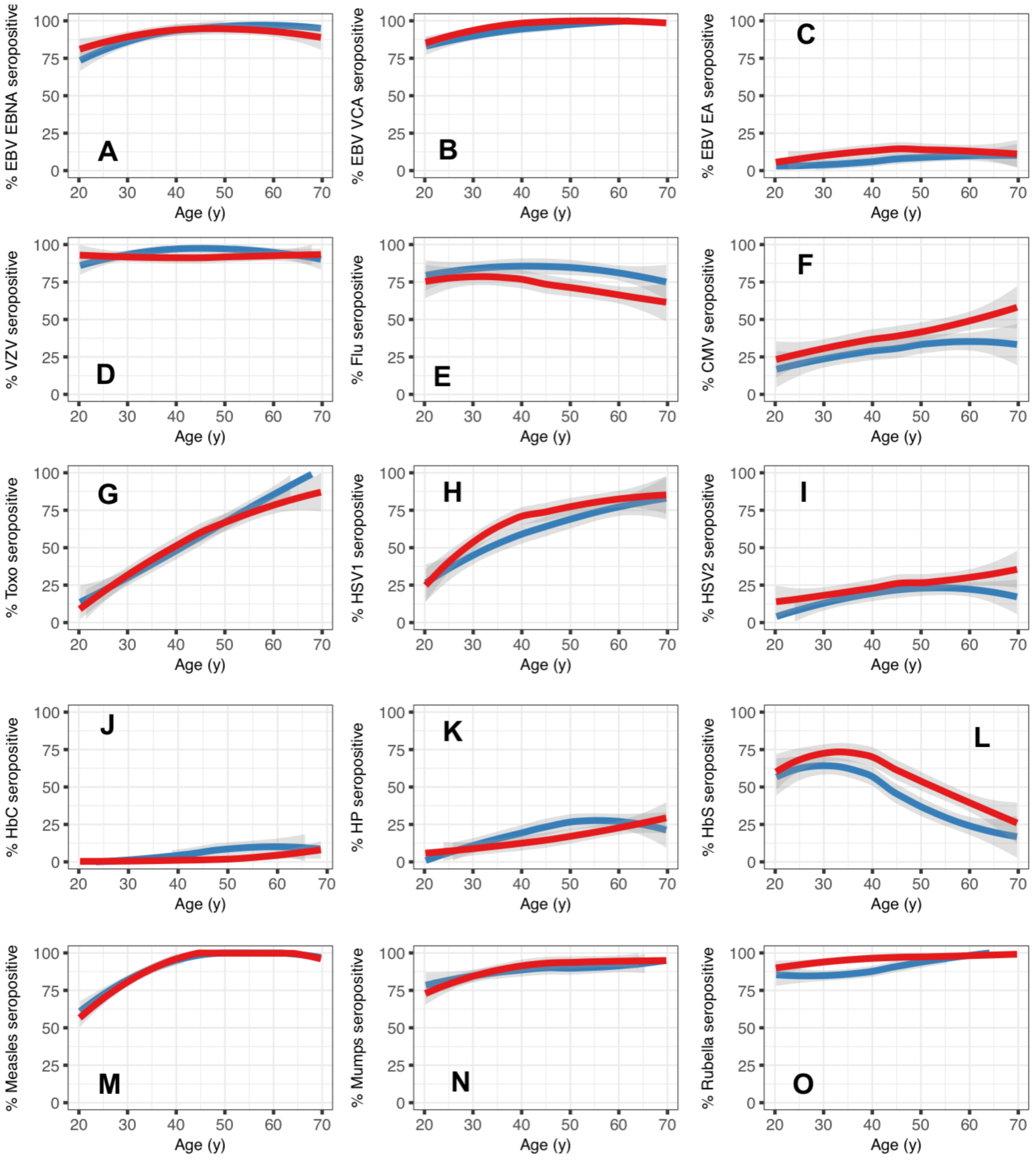
Evolution of serostatus with age and sex. (**A-O**) Odds of being seropositive for each of the 15 antigens considered in our study, as a function of age in men (blue) and women (red). Indicated *P*-values were obtained using a logistic regression with Wald test, with serostatus binary variables (seropositive, versus seronegative) as the response, and age and sex as covariates.

**Fig. S5.**
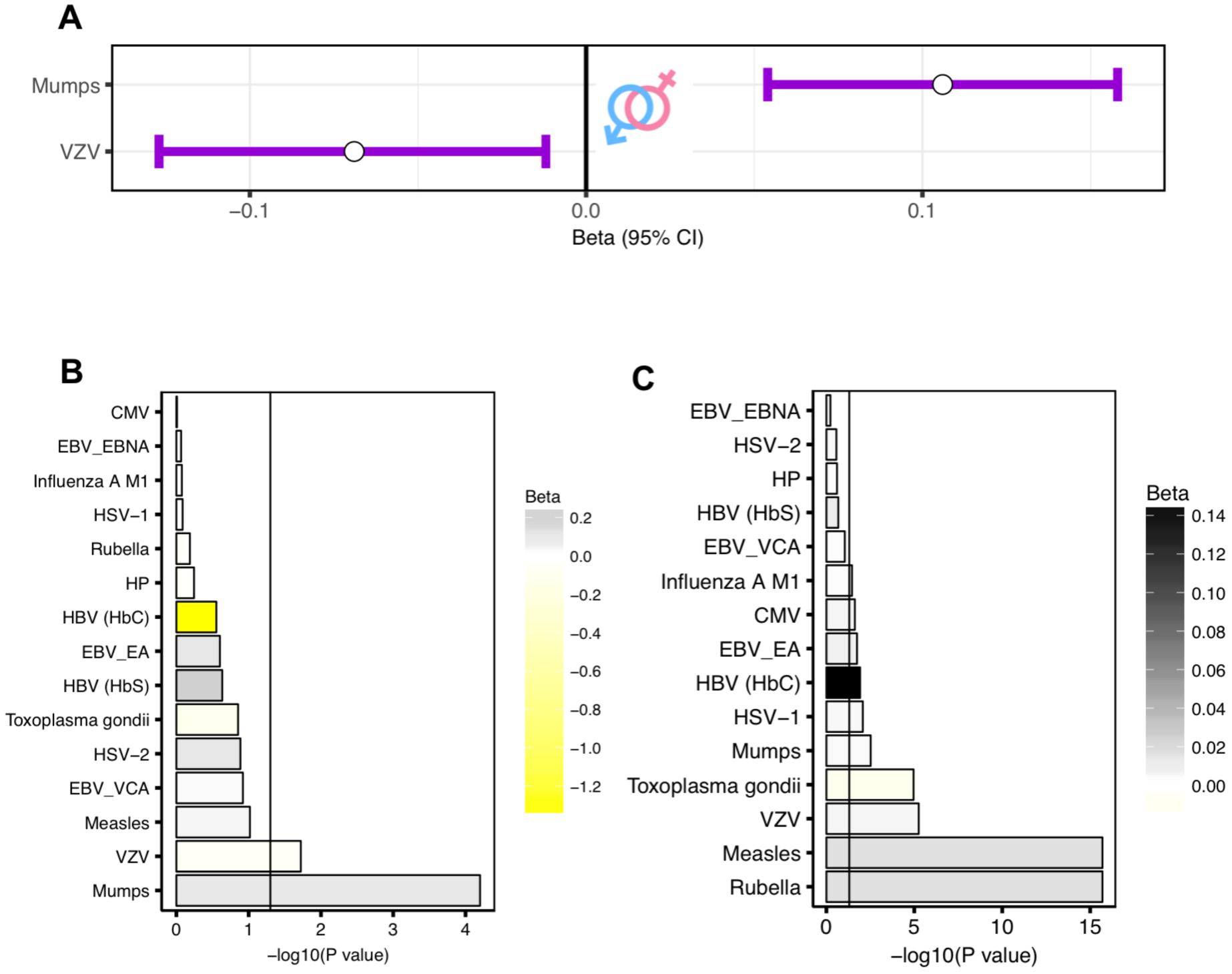
Impact of age and sex on antigen-specific IgG levels. (**A**) Effect sizes of significant associations (adjusted *P*-values (adj. *P*<0.05) between sex and Log10-transformed antigen-specific IgG levels in the 1,000 healthy individuals from the *Milieu Intérieur* cohort. Effect sizes were estimated in a linear mixed model described in (**Figure 3B**). Dots represent the mean of the beta. Lines represent the 95% confidence intervals. (**B-C**) Adjusted *P*-values (adj. *P*) of the tests of effect of age (**B**) and sex (**C**) on Log10-transformed antigen-specific IgG levels, obtained using a linear mixed model, with Log10-transformed antigen-specific IgG levels as response variables, and age and sex as treatment variables. Normalized effect sizes were color-coded. The vertical black line indicates the ‐log10 of the threshold for statistical significance (-log10(0.05) = 1.30103).

